# Dissecting *psa* locus regulation in *Yersinia pestis*

**DOI:** 10.1101/782003

**Authors:** Peng Li, Xiuran Wang, Carol Smith, Yixin Shi, Joseph T Wade, Wei Sun

**Affiliations:** Department of Immunology and Microbial Disease, Albany Medical College, Albany, NY, 12208, USA; Wadsworth Center, New York State Department of Health, Albany, NY, 12208, USA; School of Life Sciences, Arizona State University. Tempe, AZ, 85287, USA; Department of Biomedical Sciences, School of Public Health University at Albany, Rensselaer, NY 12144, USA

**Author notes:** equal author contribution (Peng Li, Xiuran Wang). corresponding author: Wei Sun,. Wade, Joseph T.

**Keywords:** PsaA (pH 6 antigen), PsaE, *Yersinia pestis*

## Abstract

The pH 6 antigen of *Yersinia pestis* is a virulence factor that is expressed in response to high temperature (37 °C) and low pH (6.0). Previous studies have implicated the PsaE and PsaF regulators in the temperature- and pH-dependent regulation of *psaA*. Here, we show that PsaE levels are themselves controlled by pH and temperature, explaining the regulation of *psaA*. We identify hundreds of binding sites for PsaE across the *Y. pestis* genome, with the majority of binding sites located in intergenic regions. However, we detect direct regulation of very few genes by PsaE, suggesting either that most binding sites are non-regulatory, or that they require additional environmental cues. We also identify the precise binding site for PsaE that is required for temperature- and pH-dependent regulation of *psaA*. Thus, our data reveal the critical function that PsaE plays in regulation of *psaA*, and suggest that PsaE may have many additional regulatory targets.

## INTRODUCTION

*Yersinia pestis* is the etiologic agent of plague, a zoonotic disease that also occurs in human populations. *Y. pestis* is one of the most lethal bacterial pathogens, and has caused several huge catastrophes in human history (Perry & Fetherston, 1997). *Y. pestis* employs a range of virulence factors that confer efficient adherence to host cells/tissues, subvert host functions, and enable it to combat host defenses. Several virulence determinants in *Y. pestis* for responsiveness to environmental cues have been identified (Cornelis, 2002, Yamashita *et al*., 2011, Lindler & Tall, 1993). Among them, PsaA (“pH 6 antigen”) was first characterized in 1961 by Ben-Efraim and coworkers, and exists in all strains of *Y. pestis*. PsaA has been reported to be synthesized only at temperatures above 34 °C and pH below 6.7 (Ben-Efraim *et al*., 1961). *Y. pestis* PsaA was reported to be a surface-exposed structure mediating agglutination of erythrocytes from several mammalian species (Bichowsky-Slomnicki & Ben-Efraim, 1963). PsaA forms a fimbrial structure with 3-5 nm diameter, and expression was reported to be induced by intracellular association with macrophages (Lindler & Tall, 1993). *Y. pestis* PsaA does not enhance adhesion to mouse macrophages, but promotes resistance to phagocytosis (Huang & Lindler, 2004). *Yersinia* PsaA is responsible for thermo-inducible adhesion to cultured mammalian cells in addition to mediating haemagglutination (Bichowsky-Slomnicki & Ben-Efraim, 1963, Felek *et al*., 2007, Galvan *et al*., 2007, Lindler & Tall, 1993, Yang *et al*., 1996, Payne *et al*., 1998, Zav’yalov *et al*., 1996, Makoveichuk *et al*., 2003).

Expression of PsaA is positively regulated by low pH combined with mammalian temperature, and by the PsaE and PsaF proteins encoded in the upstream locus (*psaEF*) (Price *et al*., 1995, Lindler *et al*., 1990, Yang & Isberg, 1997). A previous study reported that in-frame deletion of *psaE* and *psaF* greatly reduced expression of PsaA, and caused defective hemagglutination, presumably due to loss of PsaA expression (Yang & Isberg, 1997). It is unclear from previous studies how pH, temperature, and PsaE/F function to control expression of PsaA. Here, we show that PsaE directly activates transcription of *psaA* by binding to one or more DNA sites in the *psaA* upstream region. PsaE levels are controlled at the transcriptional and post-transcriptional level by pH and temperature, explaining why PsaA is expressed only at high temperature and low pH. We show that PsaE binds over 200 sites in the *Y. pestis* genome, primarily in A/T-rich intergenic regions, but directly controls expression of only two operons: *psaABC* and *psaEF*. We identify two binding sites for PsaA in the *psaA* upstream region that are required for activation by PsaE. Thus, our data reveal PsaE to be a transcription factor whose expression responds to specific environmental signals, and that binds DNA with relatively low specificity but regulates transcription in a highly specific manner.

## RESULTS

### Transcription of *Y. pestis psaA* is controlled by temperature and pH

Previous studies showed that *Y. pestis* PsaA expression is only detected in cells grown at temperatures above 36 °C and pH below 6.7 (Lindler *et al*., 1990, Ben-Efraim *et al*., 1961, Price *et al*., 1995). Using Western blotting with a PsaA-specific antibody, we confirmed this result: we were unable to detect PsaA expression from *Y. pestis* KIM6+ cells grown at 28 °C and pH 6.0 (28 °C/pH 6.0), 28 °C and pH 8.0 (28 °C/pH 8.0), and 37 °C and pH 8.0 (37 °C/pH 8.0), but we detected robust expression at 37 °C and pH 6.0 (37 °C/pH 6.0) (Figure 1A).

**Figure 1.**
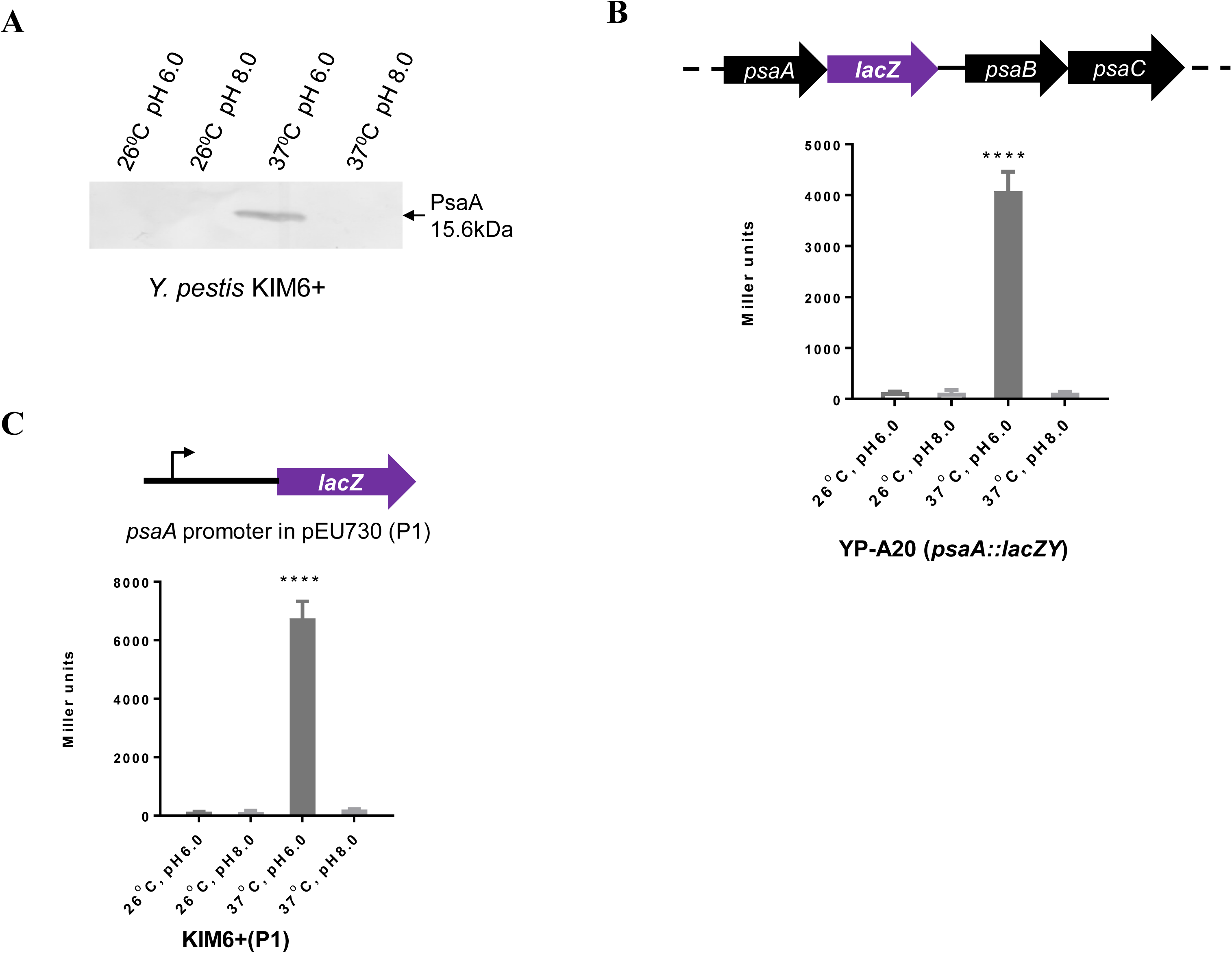
Expression of PsaA is regulated at the level of transcription by temperature and pH. (A) Western blot showing PsaA protein for wild-type KIM6+ cells grown at the indicated temperature and pH. (B) β-galactosidase of a *psaA::lacZY* fusion, for wild-type KIM6+ cells grown at the indicated temperature and pH. (C) β-galactosidase of a plasmid-borne *psaA*::*lacZ* transcriptional fusion (P1; shown in the schematic above the graph) for wild-type KIM6+ cells grown at the indicated temperature and pH. Panel A shows a representative example from three biological replicates. Data in panels B and C are from three independent assays conducted in duplicate; values shown are means, and error bars show one standard deviation. ****, *P<0.0001*; ns, not significant.

To determine whether temperature-dependent, pH-dependent regulation of PsaA occurs at the transcriptional or post-transcriptional level, we inserted the *lacZY* reporter construct downstream of the *psaA* gene in the *Y. pestis* KIM6+ chromosome. In this strain, *lacZ* expression, and hence β-galactosidase activity, is completely dependent upon transcription of the upstream *psaA*. We detected robust β-galactosidase activity for cells grown at 37 °C/pH 6.0, but not for cells grown at 28 °C/pH 6.0, 28 °C/pH 8.0, or 37 °C/pH 8.0 (Figure 1B). We conclude that temperature and pH modulate expression of *psaA* at the level of transcription, consistent with previous studies in *Y. pestis* (Price *et al*., 1995) and *Y. pseudotuberculosis* (Yang & Isberg, 1997).

To investigate the chromosomal region required for transcription regulation of *psaA*, we transcriptionally fused the full 529 bp *psaA* upstream region to a promoterless *lacZY* in a reporter plasmid (referred to P1; Table 1). We then transformed the P1 plasmid into *Y. pestis* KIM6+ and determined *lacZY* expression using a β-galactosidase assay. We detected robust β-galactosidase activity for cells grown at 37 °C/pH 6.0, but not for cells grown at 28 °C/pH 6.0, 28 °C/pH 8.0 or 37 °C/pH 8.0 (Figure 1B). We conclude that the cis-acting elements required for temperature- and pH-dependent modulation of *psaA* expression are located in the upstream region.

**Table 1.**
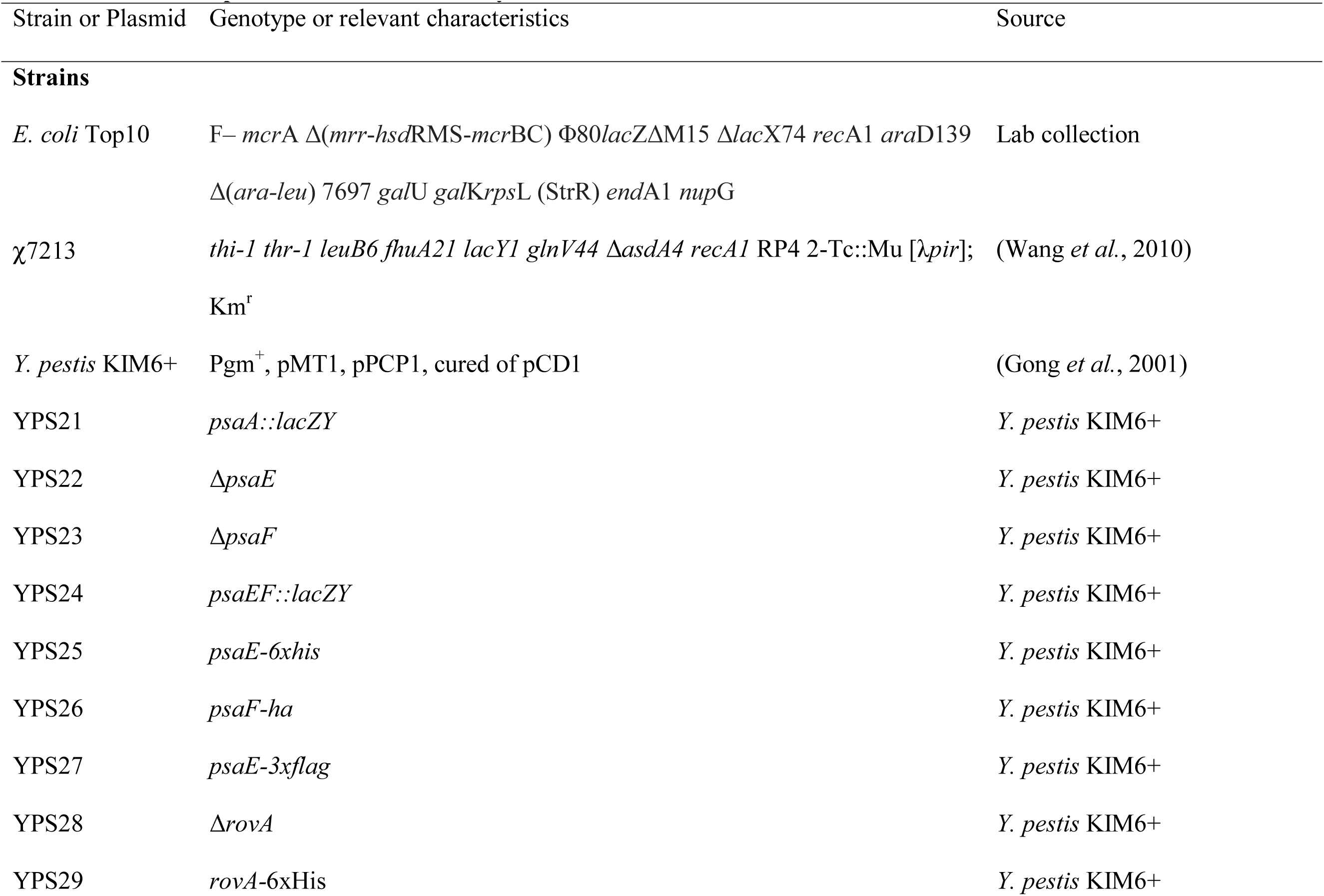
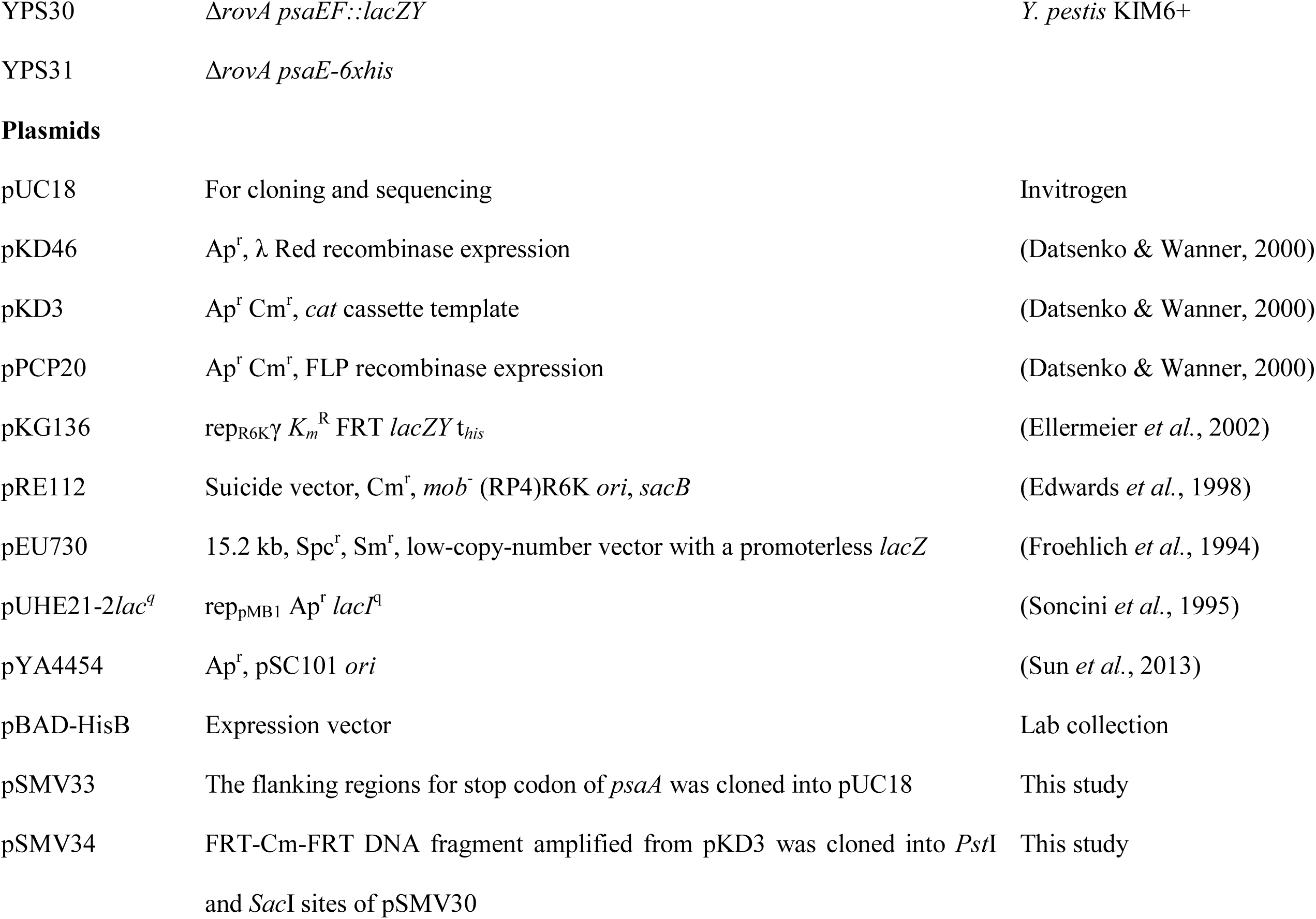
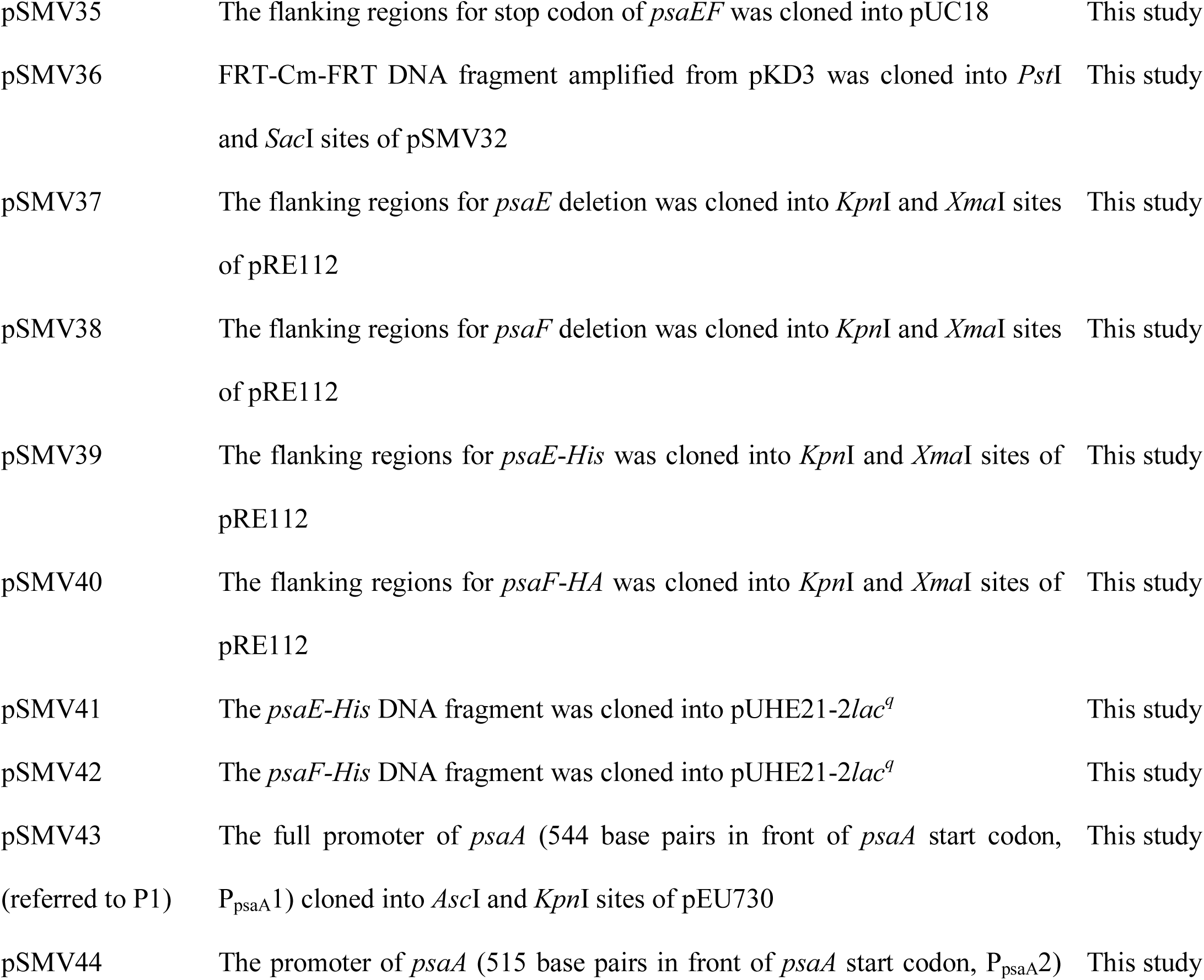
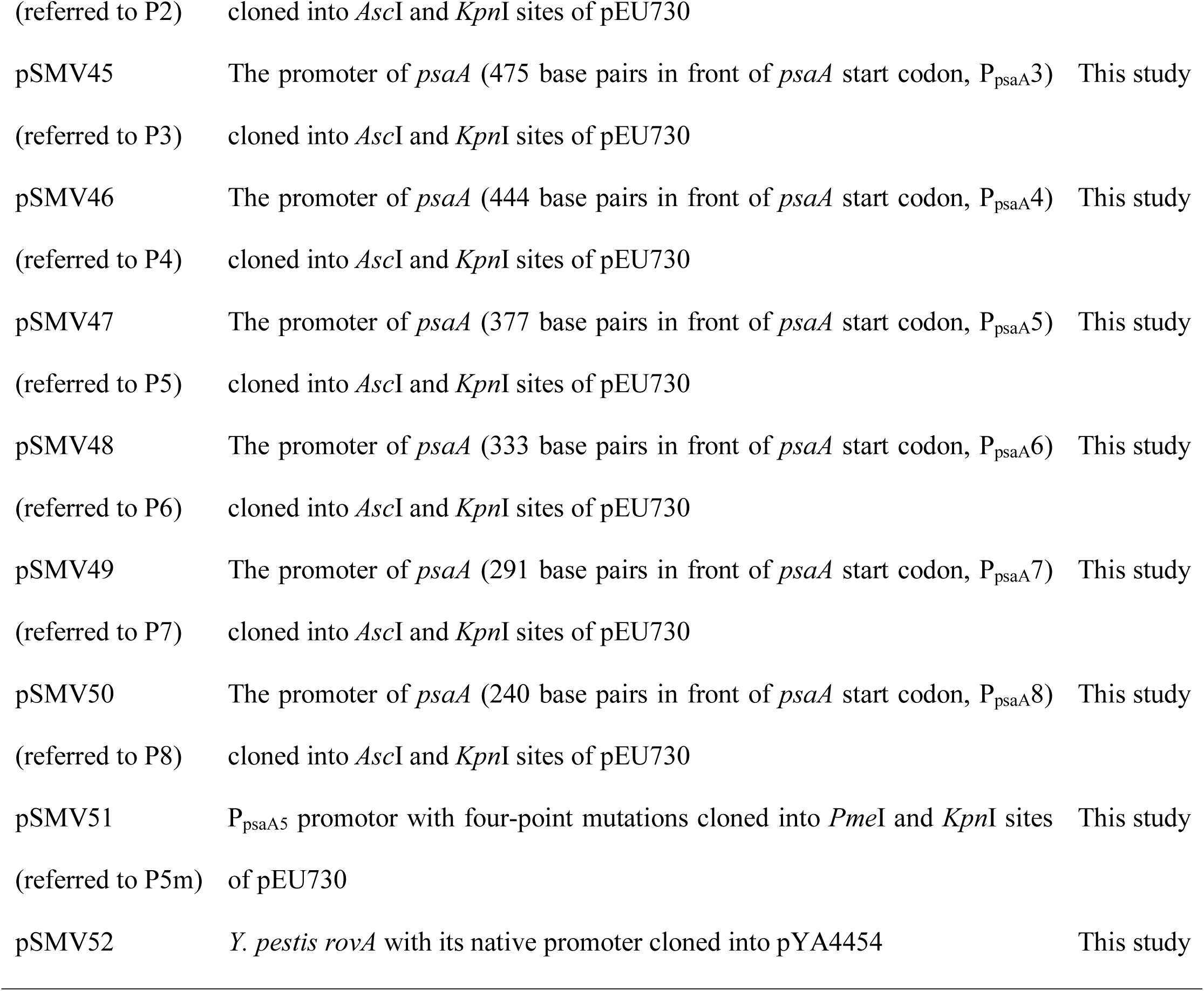
Strains and plasmids used in this study

| Strain or Plasmid | Genotype or relevant characteristics | Source |
| --- | --- | --- |
| **Strains** |  |  |
| *E. coli* Top10 | F<sup>-</sup> *mcrA* $\Delta(mrr-hsdRMS-mcrBC)$ $\Phi80lacZ\Delta M15$ $\Delta lacX74$ *recA1* *araD139*<br>$\Delta(ara-leu)$ 7697 *galU* *galK* *rpsL* (Str<sup>R</sup>) *endA1* *nupG* | Lab collection |
| χ7213 | *thi-1* *thr-1* *leuB6* *fhuA21* *lacY1* *glnV44* $\Delta asdA4$ *recA1* RP4 2-Tc::Mu [λpir];<br>Km<sup>r</sup> | (Wang *et al.*, 2010) |
| *Y. pestis* KIM6+ | Pgm<sup>+</sup>, pMT1, pPCP1, cured of pCD1 | (Gong *et al.*, 2001) |
| YPS21 | *psaA::lacZY* | *Y. pestis* KIM6+ |
| YPS22 | $\Delta psaE$ | *Y. pestis* KIM6+ |
| YPS23 | $\Delta psaF$ | *Y. pestis* KIM6+ |
| YPS24 | *psaEF::lacZY* | *Y. pestis* KIM6+ |
| YPS25 | *psaE-6xhis* | *Y. pestis* KIM6+ |
| YPS26 | *psaF-ha* | *Y. pestis* KIM6+ |
| YPS27 | *psaE-3xflag* | *Y. pestis* KIM6+ |
| YPS28 | $\Delta rovA$ | *Y. pestis* KIM6+ |
| YPS29 | *rovA-6xHis* | *Y. pestis* KIM6+ |
| YPS30 | $\Delta rovA$ *psaEF::lacZY* | *Y. pestis* KIM6+ |
| YPS31 | $\Delta rovA$ *psaE-6xhis* | |
| **Plasmids** |  |  |
| pUC18 | For cloning and sequencing | Invitrogen |
| pKD46 | Ap<sup>r</sup>, $\lambda$ Red recombinase expression | (Datsenko & Wanner, 2000) |
| pKD3 | Ap<sup>r</sup> Cm<sup>r</sup>, *cat* cassette template | (Datsenko & Wanner, 2000) |
| pPCP20 | Ap<sup>r</sup> Cm<sup>r</sup>, FLP recombinase expression | (Datsenko & Wanner, 2000) |
| pKG136 | rep<sub>R6K</sub>γ *K<sub>m</sub><sup>R</sup>* FRT *lacZY* t<sub>his</sub> | (Ellermeier *et al.*, 2002) |
| pRE112 | Suicide vector, Cm<sup>r</sup>, *mob*<sup>-</sup> (RP4)R6K *ori*, *sacB* | (Edwards *et al.*, 1998) |
| pEU730 | 15.2 kb, Spc<sup>r</sup>, Sm<sup>r</sup>, low-copy-number vector with a promoterless *lacZ* | (Froehlich *et al.*, 1994) |
| pUHE21-2*lac<sup>q</sup>* | rep<sub>pMB1</sub> Ap<sup>r</sup> *lacI<sup>q</sup>* | (Soncini *et al.*, 1995) |
| pYA4454 | Ap<sup>r</sup>, pSC101 *ori* | (Sun *et al.*, 2013) |
| pBAD-HisB | Expression vector | Lab collection |
| pSMV33 | The flanking regions for stop codon of *psaA* was cloned into pUC18 | This study |
| pSMV34 | FRT-Cm-FRT DNA fragment amplified from pKD3 was cloned into *Pst*I and *Sac*I sites of pSMV30 | This study |
| pSMV35 | The flanking regions for stop codon of *psaEF* was cloned into pUC18 | This study |
| pSMV36 | FRT-Cm-FRT DNA fragment amplified from pKD3 was cloned into *Pst*I and *Sac*I sites of pSMV32 | This study |
| pSMV37 | The flanking regions for *psaE* deletion was cloned into *Kpn*I and *Xma*I sites of pRE112 | This study |
| pSMV38 | The flanking regions for *psaF* deletion was cloned into *Kpn*I and *Xma*I sites of pRE112 | This study |
| pSMV39 | The flanking regions for *psaE-His* was cloned into *Kpn*I and *Xma*I sites of pRE112 | This study |
| pSMV40 | The flanking regions for *psaF-HA* was cloned into *Kpn*I and *Xma*I sites of pRE112 | This study |
| pSMV41 | The *psaE-His* DNA fragment was cloned into pUHE21-2*lac*<sup>q</sup> | This study |
| pSMV42 | The *psaF-His* DNA fragment was cloned into pUHE21-2*lac*<sup>q</sup> | This study |
| pSMV43<br>(referred to P1) | The full promoter of *psaA* (544 base pairs in front of *psaA* start codon, P<sub>psaA</sub>1) cloned into *Asc*I and *Kpn*I sites of pEU730 | This study |
| pSMV44 | The promoter of *psaA* (515 base pairs in front of *psaA* start codon, P<sub>psaA</sub>2) | This study |
| (referred to P2) | cloned into *AscI* and *KpnI* sites of pEU730 |  |
| pSMV45 | The promoter of *psaA* (475 base pairs in front of *psaA* start codon, P<sub>psaA</sub>3) | This study |
| (referred to P3) | cloned into *AscI* and *KpnI* sites of pEU730 |  |
| pSMV46 | The promoter of *psaA* (444 base pairs in front of *psaA* start codon, P<sub>psaA</sub>4) | This study |
| (referred to P4) | cloned into *AscI* and *KpnI* sites of pEU730 |  |
| pSMV47 | The promoter of *psaA* (377 base pairs in front of *psaA* start codon, P<sub>psaA</sub>5) | This study |
| (referred to P5) | cloned into *AscI* and *KpnI* sites of pEU730 |  |
| pSMV48 | The promoter of *psaA* (333 base pairs in front of *psaA* start codon, P<sub>psaA</sub>6) | This study |
| (referred to P6) | cloned into *AscI* and *KpnI* sites of pEU730 |  |
| pSMV49 | The promoter of *psaA* (291 base pairs in front of *psaA* start codon, P<sub>psaA</sub>7) | This study |
| (referred to P7) | cloned into *AscI* and *KpnI* sites of pEU730 |  |
| pSMV50 | The promoter of *psaA* (240 base pairs in front of *psaA* start codon, P<sub>psaA</sub>8) | This study |
| (referred to P8) | cloned into *AscI* and *KpnI* sites of pEU730 |  |
| pSMV51 | P<sub>psaA</sub>5 promotor with four-point mutations cloned into *PmeI* and *KpnI* sites | This study |
| (referred to P5m) | of pEU730 |  |
| pSMV52 | *Y. pestis rovA* with its native promoter cloned into pYA4454 | This study |

### PsaE and PsaF are required for transcription of *psaA*

Previous studies have shown that expression of *psaA* is dependent upon PsaE in *Y. pestis*, and is dependent upon PsaE and PsaF in *Y. pseudotuberculosis* (Price *et al*., 1995, Yang & Isberg, 1997, Lindler *et al*., 1990). To determine whether PsaE and PsaF regulate *Y. pestis psaA* expression via the *cis*-acting element(s) in the upstream region, we introduced the P1 *psaA*::*lacZ* reporter plasmid into wild-type *Y. pestis* KIM6+, and into Δ*psaE* and Δ*psaF* derivates. We detected robust β-galactosidase activity at 37 °C/pH 6.0, and activity was abolished in the Δ*psaE* and Δ*psaF* mutants (Figure 2A). Similarly, we detected robust expression of PsaA by Western blot in wild-type *Y. pestis* KIM6+, but not in the Δ*psaE* or Δ*psaF* mutants (Figure 2B). Expression of the *psaA*::*lacZ* reporter was restored in the Δ*psaE* and Δ*psaF* mutants respectively upon expression of respective PsaE or PsaF from an inducible plasmid (Figure 2C-D). In all cases, expression of the *psaA*::*lacZ* reporter was only detected at 37 °C/pH 6.0. We conclude that both PsaE and PsaF are required for the transcriptional induction of *Y. pestis psaA* by temperature and pH, and that PsaE/F function through *cis*-acting elements in the *psaA* upstream region.

**Figure 2.**
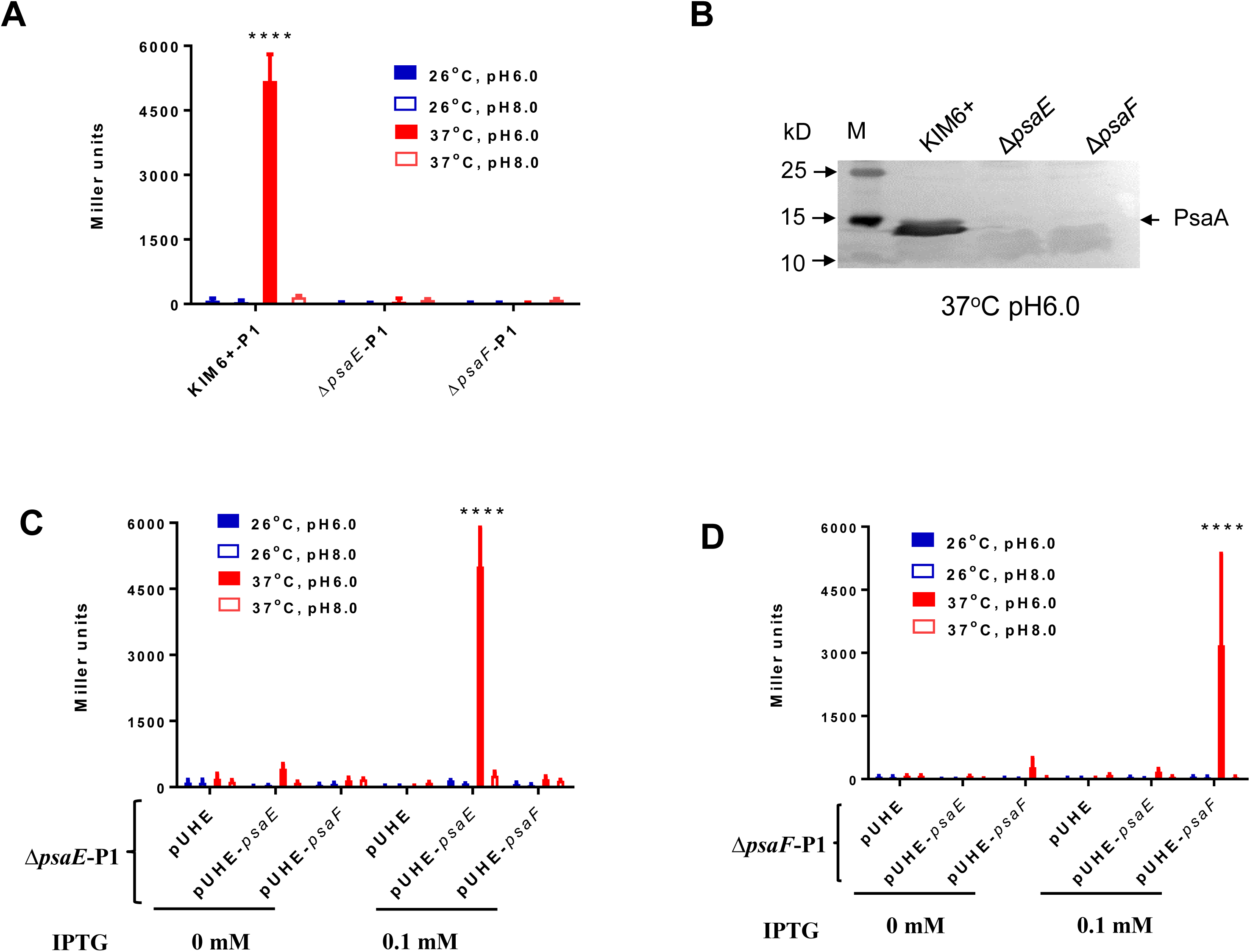
PsaE and PsaF are required for transcription activation of *psaA*. (A) β-galactosidase activity of the P1 *psaA*::*lacZ* transcriptional fusion in wild-type *Y. pestis* KIM6+, Δ*psaE* (YPS22) and Δ*psaF* (YPS23) cells grown at the indicated temperature and pH. (B) Western blot showing PsaA protein for wild-type KIM6+ cells, KIM6+ Δ*psaE*, and KIM6+ Δ*psaF* cells grown at 37 °C/pH 6.0. (C) β-galactosidase activity of the P1 *psaA*::*lacZ* transcriptional fusion in *Y. pestis* KIM6+ Δ*psaE* (YPS22) cells complemented with empty vector (pUHE), vector expressing *psaE* (pUHE-*psaE*), or vector expressing *psaF* (pUHE-*psaF*), with or without IPTG inducer, grown at the indicated temperature and pH. (D) β-galactosidase activity of the P1 *psaA*::*lacZ* transcriptional fusion in *Y. pestis* KIM6+ Δ*psaF* (YPS23) cells complemented with empty vector (pUHE), vector expressing *psaE* (pUHE-*psaE*), or vector expressing *psaF* (pUHE-*psaF*), with or without IPTG inducer, grown at the indicated temperature and pH. (D) The PsaA synthesis in the Δ*psaE* and Δ*psaF* mutants, *Y. pestis* KIM6+ as a positive control. Data are from three independent assays conducted in duplicate; values shown are means, and error bars show one standard deviation ****, *P<0.0001*; ns, not significant.

### PsaE expression is controlled transcriptionally and post-transcriptionally by temperature and pH

Since regulation of *psaA* transcription by temperature and pH is dependent upon PsaE and PsaF (Figure 2), we speculated that regulation might be due to changes in PsaE and/or PsaF levels. We inserted the *lacZY* reporter construct downstream of the *psaEF* operon in the *Y. pestis* KIM6+ chromosome. We then measured expression of the *psaEF*::*lacZY* fusion using a β-galactosidase assay. We detected β-galactosidase activity for cells grown at 28 °C/pH 6.0, 28 °C/pH 8.0, 37 °C/pH 6.0 and 37 °C/pH 8.0. However, activity was ∼3-fold higher for cells grown at 37 °C/pH 6.0 (Figure 3A). We observed a similar, but more pronounced increase in PsaE protein levels by Western blot for cells grown at 37 °C/pH 6.0 (Figure 3B).

**Figure 3.**
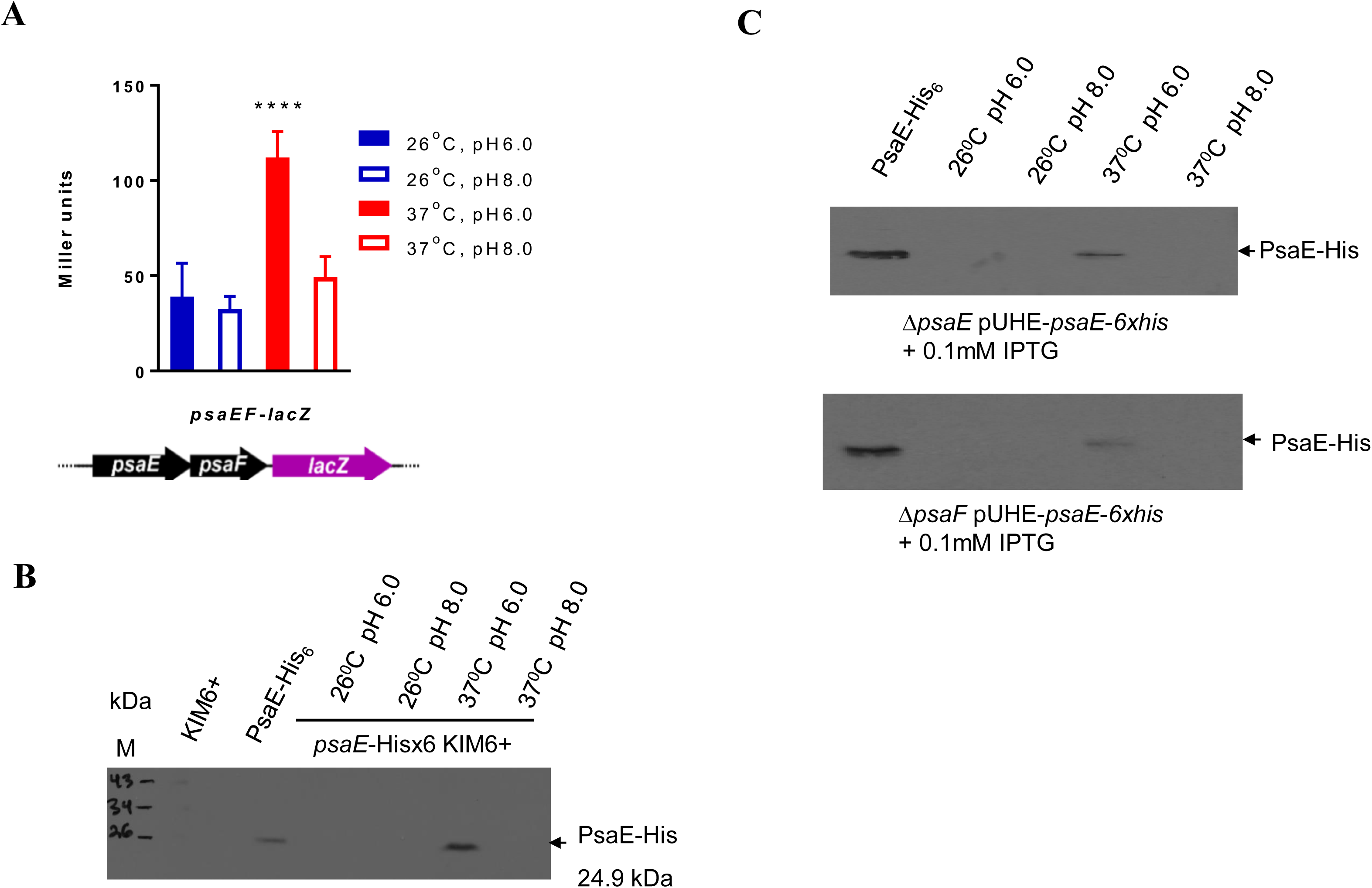
The transcription and expression of *psaEF* at four different conditions. (A) β-galactosidase activity of the *psaEF*::*lacZ* transcriptional fusion in wild-type *Y. pestis* KIM6+ cells grown at the indicated temperature and pH. (B) Western blot showing His-tagged protein expression in untagged KIM6+ cells, or KIM6+ cells expressing PsaE-His from its native locus (YPS25), grown at the indicated temperature and pH. PsaE-His overexpressed and purified from *E. coli* was run as a control. (C) Western blot showing His-tagged protein expression in KIM6+ Δ*psaE* (YPS22) or KIM6+ Δ*psaF* (YPS23) cells expressing PsaE-His from an inducible plasmid (pUHE-*psaE*-*6xHis*) in the presence of the IPTG inducer, grown at the indicated temperature and pH. PsaE-His overexpressed and purified from *E. coli* was run as a control. Data in panel A are from three independent assays conducted in duplicate; values shown are means, and error bars show one standard deviation. Panels B and C show representative examples from three biological replicates. ****, *P<0.0001*; ns, not significant.

A previous study showed that *psaE* and *psaF* transcription in *Y. pseudotuberculosis* is not subject to regulation by variations of temperature or pH (Yang & Isberg, 1997). Moreover, *Y. pestis* PsaE protein levels were undetectable in the non-inducing conditions whereas expression of the *psaEF*::*lacZY* fusion was only modestly induced by temperature and pH (compare Figure 3A and 3B). Therefore, we speculated that PsaE expression might also be controlled post-transcriptionally. To test whether PsaE expression is regulated at the level of protein production and/or stability, we overexpressed PsaE from an inducible plasmid in *Y. pestis* KIM6+ Δ*psaE* and measured PsaE protein levels by Western blot. We only detected PsaE in cells grown at 37 °C/pH 6.0 (Figure 3C), indicating that PsaE is regulated post-transcriptionally by temperature and pH. Post-transcriptional regulation of PsaE was also observed in a *Y. pestis* KIM6+ Δ*psaF* strain, indicating that regulation is not dependent upon PsaF.

### Genome-wide detection of PsaE binding and regulation

The N-terminal region of PsaE (amino acids 1-94) is predicted to bind DNA (https://www.uniprot.org/uniprot/P68588), suggesting that PsaE regulates *psaA* transcription by functioning as a DNA-binding transcription factor. To determine the PsaE regulon, we used a combination of ChIP-seq and RNA-seq to identify sites of PsaE binding and regulation on a genomic scale. ChIP-seq identifies binding sites for DNA-binding proteins, but does not provide information on whether binding DNA is associated to transcription regulation. RNA-seq can identify changes in RNA levels that are dependent upon a transcription factor, but cannot be used to determine whether instances of regulation are direct or indirect. By combining ChIP-seq and RNA-seq, it is possible to identify sites of direct regulation.

We constructed PsaE-FLAG and PsaE-His tagged strains of *Y. pestis* KIM6+, with the tags introduced at the native *psaE* locus. Insertion of the tags did not affect PsaA expression levels (Fig. S1). We then performed ChIP-seq for each of the tagged strains. We detected 218 sites of PsaE enrichment across the chromosome, with the pattern of enrichment being very similar for each of the two tagged strains (Figure 4A). No equivalent enrichment was detected in control ChIP-seq experiments using an untagged strain (Figure 4A). One of the most highly enriched regions was identified upstream of *psaE*, with strong enrichment also observed nearby, upstream of *psaE* (Figure 4B). We identified a highly enriched sequence motif within the 218 PsaE-bound regions (Figure 4B; Supplementary Table 1), with an instance of the motif being found for each of the 218 regions. The sequence motif was significantly centrally enriched with respect to the peaks of ChIP-seq signal (*p* = 2.9e^-11^), strongly suggesting that the motif corresponds to the PsaE binding sequence.

**Figure 4.**
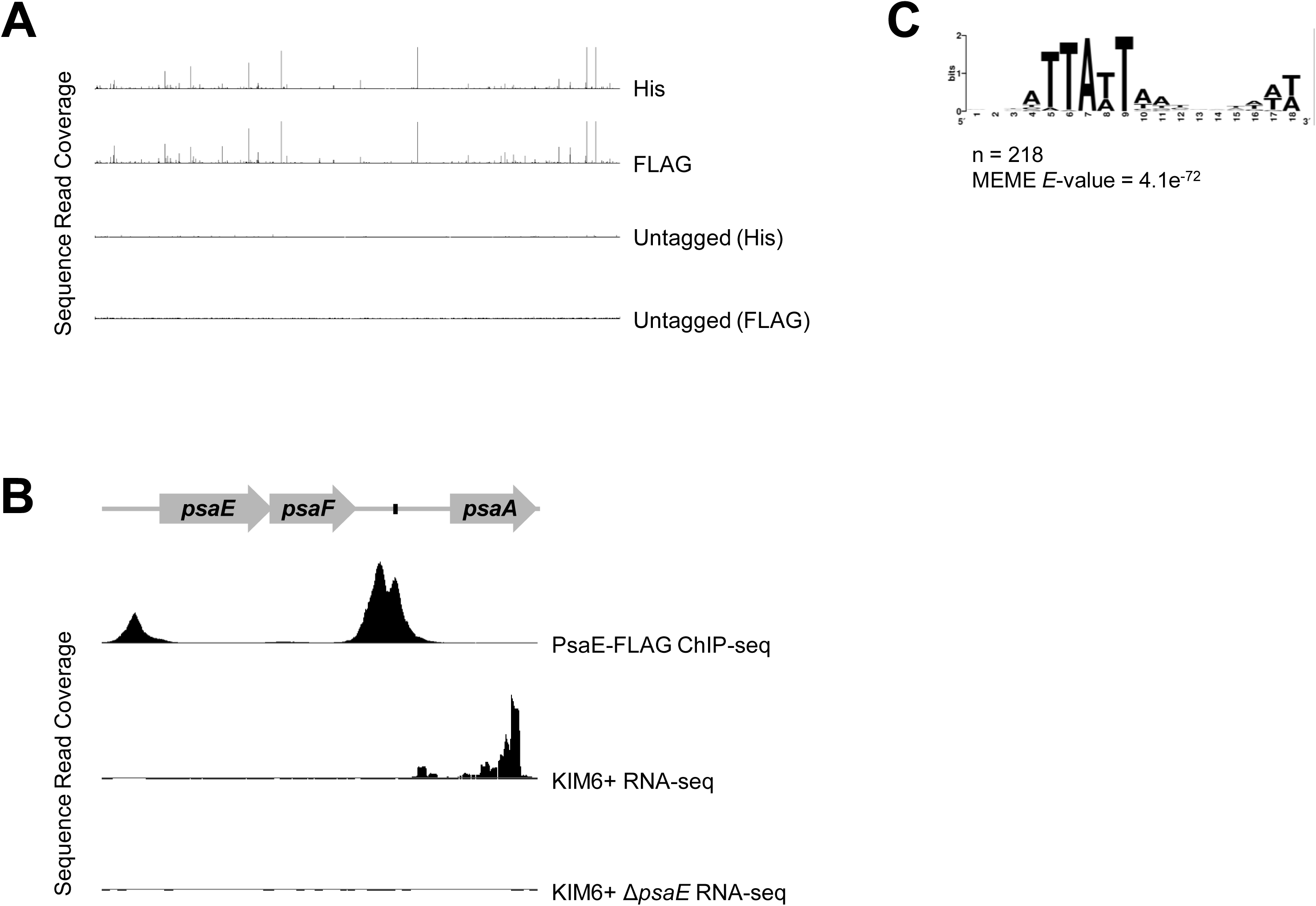
ChIP-seq of PsaE identifies 218 binding sites that are strongly associated with a sequence motif. (A) Overview of the ChIP-seq data for PsaE-His6, Psa-FLAG, and untagged controls. For the untagged control datasets, the antibody used in the ChIP-seq is indicated in parentheses. Graphs show sequence read coverage on the plus strand from ChIP-enriched samples (y-axis) across the entire *Y. pestis* chromosome (x-axis). Peaks correspond to PsaE binding sites. Values on the y-axis are capped at one tenth the level of the most enriched region in the PsaE-His dataset, and the maximum value shown in each plot is normalized between datasets based on the total number of sequence reads that align to the reference genome. (B) Zoomed-in view of the *psa* locus. ChIP-seq data are shown for the PsaE-FLAG dataset, and RNA-seq data are shown for wild-type KIM6+ and KIM6+ Δ*psaE* strains. Gene positions are indicated in the schematic above the graphs. The black box indicates the location of the putative PsaE binding site mutated in the P5m construct (see Figure 6). (C) A strongly enriched motif was associated with PsaE ChIP-seq peaks.

Most ChIP-seq studies of bacterial DNA-binding transcription factors identify <50 sites. However, in some cases, transcription factors have been observed to bind at hundreds of DNA sites, as is the case for PsaE (Galagan *et al*., 2013b). In cases where transcription factors bind >100 sites, most sites are usually located within genes, suggesting a random distribution of non-regulatory sites across the chromosome (Galagan *et al*., 2013b). Strikingly, 54% of PsaE-bound sites fall in intergenic regions, a far higher fraction than the 17% of the genome that is intergenic. Thus, PsaE binding is strongly enriched for intergenic regions. Moreover, PsaE binding is strongly associated with A/T-rich sequence, with only four of the 218 PsaE-bound regions having an A/T content below 50%; the median A/T content of PsaE-bound regions is 67.3%. By contrast, the A/T content of the *Y. pestis* KIM6+ chromosome is 52.3%. Consistent with PsaE binding in regions of high A/T content, the PsaE binding site motif is strongly A/T-rich (Figure 4B).

We next determined which genes are regulated by PsaE by using RNA-seq to compare RNA levels genome-wide in wild-type *Y. pestis* KIM6+ and a Δ*psaE* mutant. Thus, we identified 127 differentially regulated genes between the two strains (Figure 5; Supplementary Table 2). The most strongly differentially regulated genes were *psaA*, *psaB*, *psaC*, and *psaF* (Figure 5). A previous study suggested that *psaA* is transcribed as a monocistronic RNA (Price *et al*., 1995). Although we observed PsaE-dependent regulation of the two genes downstream of *psaA* (i.e. *psaB* and *psaC*), the absolute RNA level for *psaA* is ∼1,000 times higher than that for *psaB* and *psaC*, strongly suggesting that the observed regulation of *psaB* and *psaC* by PsaE is due to low-level read-through of the *psaA* transcription terminator. Of the remaining 123 PsaE-regulated genes, only five are located within a −100 to +500 bp window relative to PsaE-bound regions identified by ChIP-seq (Figure 5; Supplementary Table 2), and these genes are associated with relatively weak PsaE binding sites, as determined by the level of ChIP-seq enrichment. Given the large number of PsaE binding sites identified by ChIP-seq, we estimate that ∼4 differentially expressed genes would be found within a −100 to +500 bp window relative to PsaE-bound region by chance. We conclude that PsaE directly regulates very few genes under the growth conditions tested, despite the large number of intergenic binding sites.

**Figure 5.**
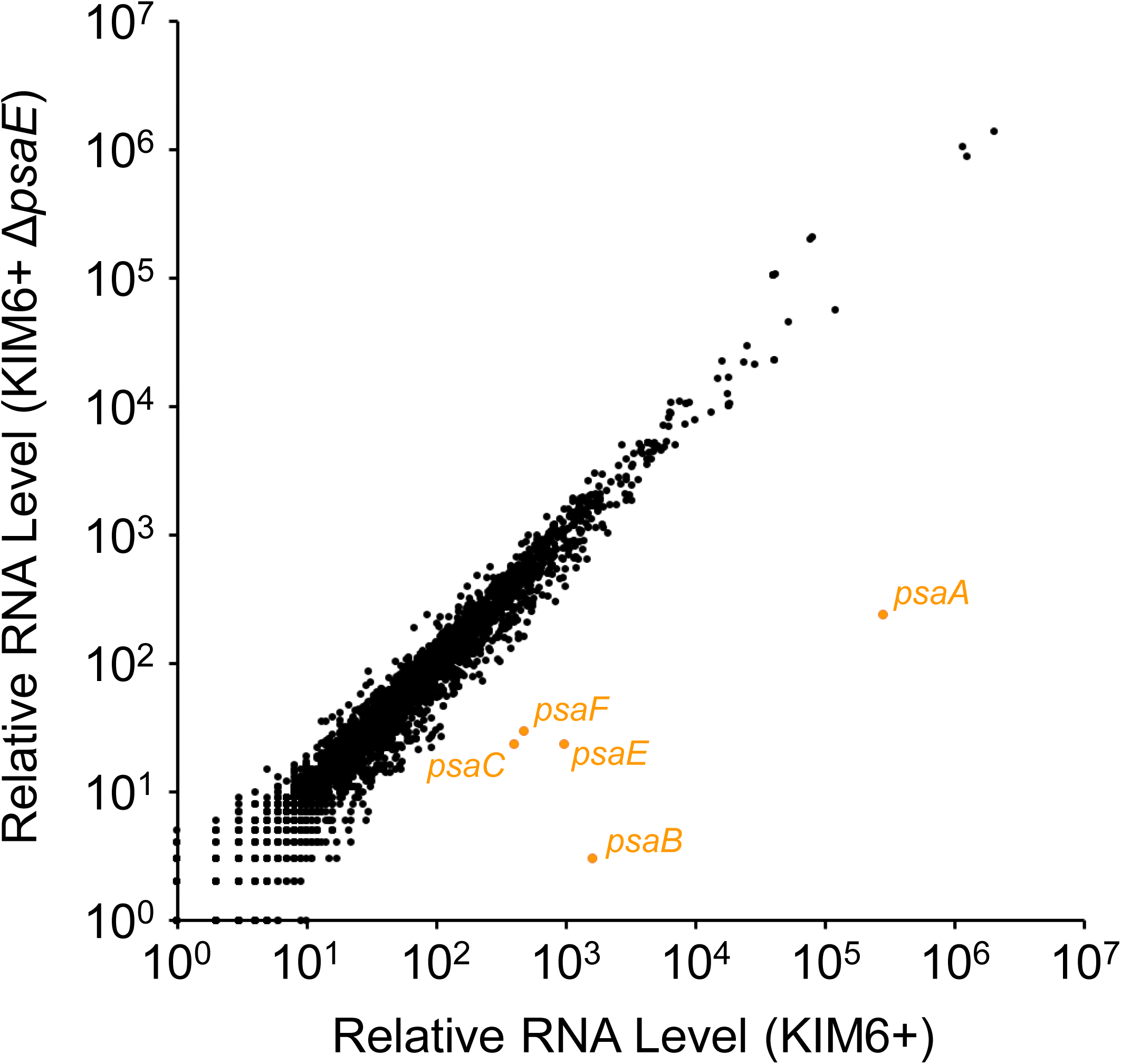
RNA-seq of wild-type and Δ*psaE* cells reveals regulation of few transcripts. Scatter-plot showing relative RNA levels for all genes, as determined by RNA-seq, for wild-type cells (x-axis) and Δ*psaE* mutant cells (y-axis). The *psaA*, *psaB*, *psaC*, *psaE* and *psaF* genes are shown in orange.

### Identifying the site of PsaE binding in *psaA* upstream region

Our data suggest that PsaE regulates transcription of *psaA* using a *cis*-acting element in the *psaA* upstream region. Given that we identified binding of PsaE upstream of *psaA*, we speculated that one or more PsaE binding site in the *psaA* upstream region is required for regulation by PsaE. There are several good matches to the PsaE binding motif in the *psaA* upstream region. Consistent with this, the profile of ChIP-seq enrichment suggests the presence of at least two PsaE binding sites (Figure 4B). To begin to map the site of PsaE binding, we made derivatives of the P1 *psaA*::*lacZ* reporter plasmid with progressive truncations of the start of the *psaA* upstream region. We refer to these reporter plasmids as P2, P3, P4, P5, P6, P7 and P8 (Figure 6A). We introduced these plasmids into *Y. pestis* KIM6+ and measured β-galactosidase activity in cells grown at 28 °C/pH 6.0, 28 °C/pH 8.0, 37 °C/pH 6.0 or 37 °C/pH 8.0. For all reporter fusions, no activity was observed under non-inducing conditions (i.e. 28 °C/pH 6.0, 28 °C/pH 8.0 and 37 °C/pH 8.0; Figure 6B). We detected robust β-galactosidase activity under inducing conditions (i.e. 37 °C/pH 6.0) for P1-P6, but not for P7 or P8 (Figure 6B). These data strongly suggest that the key binding site(s) for PsaE are located in the region from −333 to −292 relative to the start codon of *psaA*. Consistent with this idea, the region from −333 to −292 contains two strong matches to the PsaA binding motif identified from ChIP-seq data (Figure 6A), and the profile of ChIP-seq enrichment matches the position of these sequences (Figure 4B).

**Figure 6.**
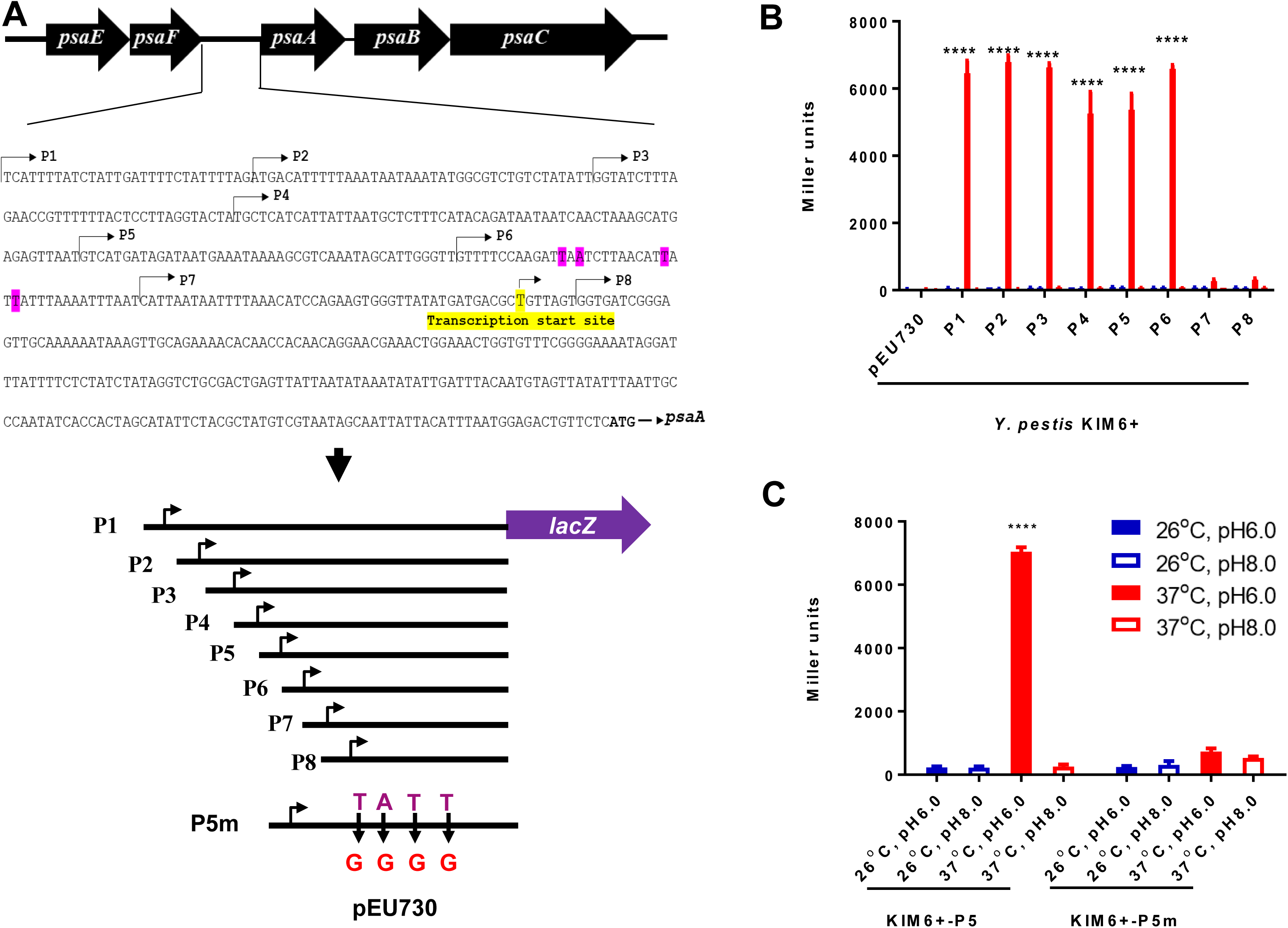
Mapping the PsaE binding region of the *psaA* promoter. (A) Schematic representation of constructs used for analyzing the sequence requirements for transcription activation of *psaA* by PsaE. Bent arrows indicate truncations of the P1 reporter construct (P2-8). Pink-highlighted bases indicate predicted PsaE binding sites that were mutated in the P5m construct. The yellow highlighted base indicates the transcription start site. (B) β-galactosidase activity of the P1-8 constructs or empty vector (pEU730) in wild-type KIM6+ cells grown at the indicated temperature and pH. (C) β-galactosidase activity of the P5 and P5m constructs in wild-type KIM6+ cells grown at the indicated temperature and pH. Data are from three independent assays conducted in duplicate; values shown are means, and error bars show one standard deviation ****, *P<0.0001*; ns, not significant. ****, *P<0.0001*; ns, not significant.

To test whether the predicted PsaE binding sites are required for transcription activation of *psaA* by PsaE, we modified the P5 plasmid by introducing two substitutions in each putative site that alter key positions as predicted from the binding site motif (compare Figure 6 and Figure 4C). We refer to this mutated plasmid as P5m. We then introduced the P5 and P5m plasmids into *Y. pestis* KIM6+ and measured β-galactosidase activity in cells grown at 28 °C/pH 6.0, 28 °C/pH 8.0, 37 °C/pH 6.0 or 37 °C/pH 8.0. Expression of the *psaA*::*lacZ* fusion under inducing conditions was abolished in the P5m mutant construct (Figure 6C). We conclude PsaE binds to one or both of the putative sites in the region from −333 to −292, and that binding to one or both of these sites is critical for transcription activation of *psaA*.

## DISCUSSION

Our data support a model in which PsaE activates transcription of *psaA* by binding to one or more DNA sites in the *psaA* upstream region. Moreover, we have shown that PsaE levels are controlled post-transcriptionally by temperature and pH, explaining why *psaA* transcription is activated at 37 °C/pH 6.0. During the course of this work, Quinn *et al*. (Quinn *et al*., 2019) also investigated the mechanism of *psaA* regulation. Consistent with our study, Quinn *et al*. showed that PsaE and PsaF are transcriptionally and translationally regulated by temperature, and post-translationally regulated by pH. However, in contrast to our study, Quinn et al. did not observe transcriptional regulation of *psaEF* by pH (Figure 3A). Interestingly, a previous study of *psaA* regulation in *Y. pseudotuberculosis* showed that *psaEF* transcription is not regulated by temperature or pH (Yang & Isberg, 1997). Thus, the mechanisms by which temperature and pH control PsaE activity may differ between species. Nonetheless, control of PsaE activity by temperature and pH appears to be a widespread phenomenon, since a *psaA* homologue in *Yersinia enterocolitica*, *myfA*, is also transcriptionally activated by high temperature and low pH, and requires homologues of *psaE* and *psaF* for maximal expression (Iriarte & Cornelis, 1995).

PsaE has an unusually large number of binding sites across entire genome (218 putative binding sites). For comparison, ChIP-seq data for CRP in Enterotoxigenic *E. coli* identified 111 sites, and CRP is considered to be a global regulator with one of the largest regulons of any *E. coli* transcription factor (Haycocks *et al*., 2015). Some bacterial transcription factors bind large numbers of sites that appear to lack regulatory activity (Galagan *et al*., 2013a, Wade *et al*., 2007). However, these sites are typically distributed proportionally between intergenic and genic locations, which means that most fall within genes, since bacterial genomes are composed mainly of genic sequence. By contrast, PsaE sites are highly enriched for intergenic regions (Binomial test *p* < 1e^-15^), strongly suggesting that PsaE regulates transcription of the downstream genes and may be a global regulator, directly controlling the transcription of hundreds of genes. However, we observed strong PsaE-dependent regulation of only two operons, *psaABC* and *psaEF*; the proximity of a handful of weakly PsaE-regulated to PsaE binding sites is likely coincidental. Our data are consistent with PsaE/F autoregulation, although reduced *psaF* expression in the Δ*psaE* mutant could be due to the altered upstream sequence for *psaF*, especially since this region has been implicated in controlling PsaF translation (Quinn *et al*., 2019).

One possible explanation for why PsaE binds so many intragenic sites but is associated with so little detectable regulation is that regulation may require additional factors that are inactive under the growth conditions we used. Consistent with this idea, several other transcription factors have been implicated in controlling expression of *psaA*, including RovA (Cathelyn *et al*., 2006, Cathelyn *et al*., 2007, Zhang *et al*., 2013), CpxR (Liu *et al*., 2011), PhoP (Zhang *et al*., 2013) and Fur (Zhou et al., 2006). Intriguingly, several PsaE binding sites, including the site with the most enrichment in the ChIP-seq data, are upstream of genes associated with iron acquisition: *feoABC*, *yiuA*, and *ftn*, suggesting that PsaE may collaborate with the same transcription factor to regulate these genes.

The very high A/T content of sequences surrounding PsaE binding sites suggests a connection to the nucleoid-associated protein H-NS. A/T-rich regions of γ-proteobacterial genomes are very strongly associated with transcriptional silencing by H-NS (Navarre *et al*., 2006, Singh *et al*., 2014, Lucchini *et al*., 2006, Grainger *et al*., 2006, Oshima *et al*., 2006), and H-NS has a well-established role in selectively repressing virulence genes in *Yersinia* (Stoebel et al., 2008). Many virulence-associated transcription factors in a wide range of bacterial species activate transcription by binding in H-NS-bound genomic regions and counteracting the repressive effects of H-NS (Stoebel *et al*., 2008, Will *et al*., 2014). We suggest that many of the intergenic PsaE binding sites are associated with regulation of the downstream genes under conditions where H-NS has been locally or globally destabilized. In such a model, binding of H-NS under most growth conditions would prevent transcription activation by PsaE.

An alternative possibility to explain the lack of detected regulation by PsaE is that the majority of PsaE binding sites are non-regulatory, and that regulation by PsaA requires strict spacing requirements relative to promoter sequences. The *psaA* transcription start site has been mapped (Zhang *et al*., 2013), and our data (Figure 6) indicate that PsaE binds ∼50 bp upstream of this site. This is consistent with the location of activator proteins relative to the binding site of RNA polymerase. Many activator proteins need to be precisely positioned on the DNA relative to promoter sequences to be effective (Lee *et al*., 2012). Hence, it is possible that the majority of PsaE sites are not suitably positioned to activate transcription. However, this does not explain why so many PsaE sites are located in intergenic regions.

In conclusion, we have characterized the regulation of *Y. pestis psaA* by temperature and pH, revealing a key role for PsaE binding to one or more promoter-proximal sites. Several key unanswered questions remain. First, what is the role of PsaA during infection? Second, how does pH impact the stability of PsaE and PsaF? Third, are there additional PsaE-regulated genes that are activated in a condition-specific manner?

## MATERIALS AND METHODS

### Bacterial Strains and Growth Conditions

All strains used in this study and their sources are listed in Table 1. Conditions and media used for growing strains of *Y. pestis* and *E. coli* were described previously (Sun *et al*., 2008). For determination of PsaA expression at various growth conditions, *Y. pestis* was grown in Heart Infusion broth (Difco) adjusted to either pH 8.0 or pH 6.0 with NaOH or HCl, respectively. After autoclaving, 2.5 mM CaCl_2_ and 0.2% (wt/vol) xylose were added (Lindler *et al*., 1990). Chloramphenicol (Cm), Spectinomycin (Sp), and ampicillin (Ap) were added to the media at final concentrations of 20, 50, and 100 μg/ml, respectively.

### Plasmid Construction

All plasmids used in this study are described in Table 1. Primers used for PCR amplification are listed in in Supplementary Table 3. DNA manipulations were performed following standard methods (Sambrook, 1989), and all constructed plasmids were confirmed by DNA sequencing.

### Strain construction

To construct *Y. pestis* mutant strains, the corresponding suicide plasmid was conjugationally transferred from *E. coli* χ7213 (Wang *et al*., 2010) to the *Y. pestis* strain. Single-crossover insertion strains were isolated on TBA agar plates containing Cm. Loss of the suicide vector sequence after the second recombination between homologous regions (i.e., allelic exchange) was selected by using the SacB-based sucrose sensitivity counter-selection system (Sun *et al*., 2008). A *lacZY* gene cassette was integrated in the deleted chromosomal *psaA* or *psaEF* location using plasmid pKG136, which was inserted into the FLP recombination target sequence generated after the FRT-Cm^R^-FRT cassette was removed using plasmid pCP20 (Ellermeier *et al*., 2002, Song *et al*., 2008).

### Western blotting

Bacteria grown at different conditions were pelleted and boiled in gel loading dye, separated by sodium dodecyl sulfate-polyacrylamide gel electrophoresis (SDS-PAGE) with 12.5% polyacrylamide. Proteins were transferred to nitrocellulose membranes for Western blot analysis. Loading was normalized based on optical density (OD_600_) of bacterial culture. Membranes were blocked with 5% skimmed milk in PBS, incubated with primary antibody, specifically rabbit polyclonal anti-PsaA (1:10,000) and mAb anti-His (1:2,000) to probe for PsaA and PsaE-His_6_, respectively. Membranes were then incubated with goat anti-rabbit immunoglobulin G (IgG) alkaline phosphatase-conjugated (1:10,000) or goat anti-mouse IgG horseradish peroxidase (HRP)-conjugated (1:10,000) secondary antibody (Sigma, St. Louis, MO). Immunoreactive bands were detected by the addition of NBT/BCIP (Sigma, St. Louis, MO) and chemiluminescent substrates (Bio-Rad, CA).

### **β**-Galactosidase Assays

β-galactosidase assays (Miller, 1972) were carried out in triplicate, and activity was determined using a SpectraMax. 340PC plate reader (Molecular Device). The data correspond to three independent assays conducted in duplicate, and values shown are the mean, with error bars indicating one standard deviation.

### ChIP-seq

Briefly, KIM6+ and YPS24 (*psaE*-His) were grown in HIB at 37 °C, pH6 to an OD_600_ of 0.6 to 0.8. Sonicated lysates were prepared as previously described (Bonocora & Wade, 2015). ChIP-seq libraries were prepared as previously described (Bonocora & Wade, 2015), using 2 µg anti-His tag monoclonal antibody and 25 µL Protein G Sepharose slurry, or 2 µL M2 anti-FLAG antibody (Sigma) and 25 µL Protein A Sepharose slurry. Sequencing was performed using an Illumina Next-Seq Instrument (Wadsworth Center Applied Genomic Technologies Core). Sequence reads were aligned to the *Y. pestis* KIM6+ genome sequence using the CLC Genomics Workbench (version 9). Statistically significant (false discover rate of 0.01) enriched regions were identified as previously described (Fitzgerald *et al*., 2014), treating the PsaE-His and PsaE-FLAG datasets as independent replicates.

### ChIP-seq motif analysis

We extracted 101 bp DNA sequence centered around each ChIP-seq peak. We merged ChIP-seq peaks within 100 bp of each other. We used MEME (version 5.0.5; default settings) to identify an enriched sequence motif (Bailey & Elkan, 1994). The position of motifs identified by MEME was compared to the ChIP-seq peak center position using Centrimo (Bailey & Machanick, 2012) (version 5.0.5; default settings).

### RNA-seq

RNA-seq was performed in strains KIM6+ and YPS21 (Δ*psaE*). Bacteria cultured overnight in pH 8.0 of HIB media at 28 °C were collected and reinoculated in the pH 6 of HIB media and cultured at 37 °C for 4 h. One ml cells were pelleted in a microcentrifuge for 1 min at full speed and washed once with Tris-buffered saline. RNA was purified from bacterial cells using the Quick-RNA™ Miniprep Plus Kit (Zymo Research). Duplicate samples were prepared from independent biological replicates for each condition/strain. 6 ug RNA was treated with 3 units of Turbo DNase I (Invitrogen) for 45 min at 37 °C. RNA was then acid phenol extracted and ethanol precipitated. Ribosomal RNA was removed using the RiboZero kit (Illumina). RNA-seq libraries were prepared using the ScriptSeq 2.0 Kit (Illumina). Sequencing was performed using an Illumina Next-Seq instrument (Wadsworth Center Applied Genomic Technologies Core). Differential RNA expression analysis was performed using Rockhopper (version 2.03) with default parameters (McClure *et al*., 2013). Differences in RNA levels were considered to indicate regulation for genes with false-discovery-rate (*q*) values of ≤0.01 and fold-change values ≥2.

### Statistical analyses

Statistical analyses of data among groups were evaluated by two-way ANOVA and Tukey’s multiple-comparison test. Data were analyzed using GraphPad PRISM 8.0 software. Data represented as mean value ± standard deviation.

## DATA AVAILABILITY

Raw ChIP-seq and RNA-seq data are available from EBI ArrayExpress with accession numbers E-MTAB-8369 and E-MTAB-8370.

## ACKNOWLEDGEMENTS

We thank Dr. Roy Curtiss III for providing *E. coli* and *Y. pestis* strains, and different plasmids in this study. We thank the Wadsworth Center Applied Genomic Technologies Core Facility for DNA sequencing. We thank the Wadsworth Center Tissue Culture and Media Core Facility and Glassware Facility for technical support. This work was primarily supported by Albany Medical College start-up fund and partially supported by National Institutes of Health grant R01AI125623 to WS. This work was also supported by National Institutes of Health grant R01GM114812 to JTW.

## AUTHOR CONTRIBUTIONS

Conceived and designed the experiments: WS, YXS, JTW. Performed the experiments: PL, XRW, CS, WS. Analyzed the data: WS, YXS, CS, JTW. Contributed reagents/materials/analysis tools: WS, YXS, JTW. Wrote the paper: WS, YXS, JTW.

