## Supplementary material for "Dissecting *psa* locus regulation in *Yersinia pestis*": Figure S1

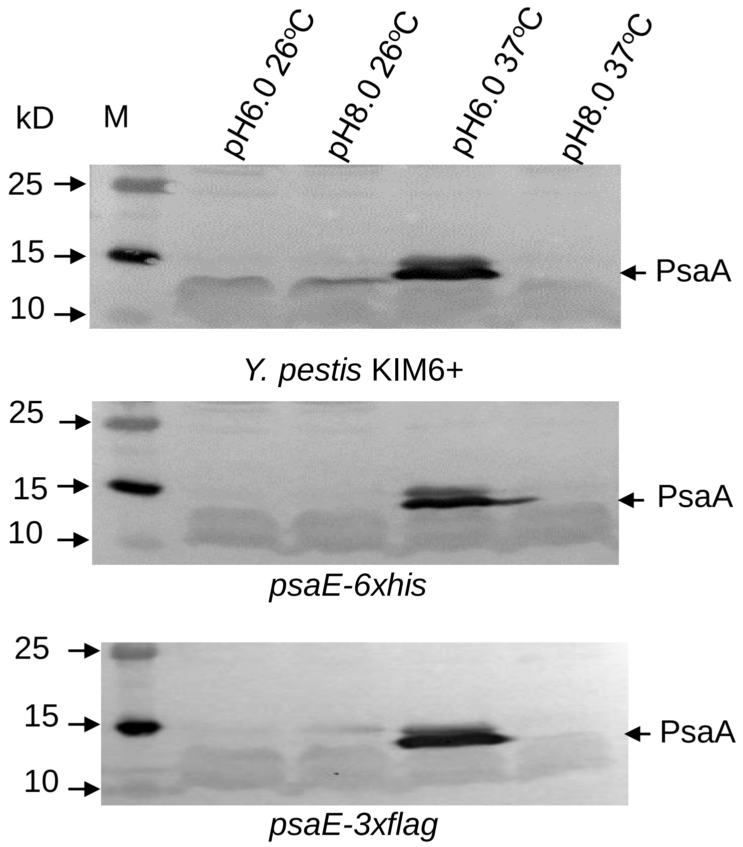


**Figure S1. PsaA protein levels in different strains and growth conditions.** PsaA protein was detected using Western blotting for *psaE*-6xHis, *psaE*-3xFLAG and wild-type *Y. pestis* KIM6+ strains grown under four different conditions (pH 6/26 °C, pH 8/26 °C, pH 6/37 °C and pH 8/37 °C).
