## Supplementary Methods for "Dissecting *psa* locus regulation in *Yersinia pestis*"

### Supplementary Materials and Methods

#### Plasmid Constructions

For construction of the *psaA::lacZY* insertion, primer sets of PsaA-Lac1/PsaA-Lac2 and PsaA-Lac3/PsaA-Lac4 were used for amplifying the upstream and the downstream sequences of the *psaA* stop codon, respectively. Complementary region in primers PsaA-Lac2 and PsaA-Lac3 contained *Pst*I and *Sac*I restriction sites. The flanking region of the *psaA* stop codon were fused by overlapping PCR using primers PsaA-Lac1and PsaA-Lac4. The resulting PCR fragments were constructed into *Eco*RI and *Hind*III sites of pUC18 to generate the plasmid pSMV33. Then, the FRT-Cm-FRT DNA fragments amplified from pKD3 was cloned into *Pst*I and *Sac*I sites of pSMV33 to generate pSMV34. For construction of the *psaEF::lacZY* insertion, primer sets of PsaEF-Lac1/PsaEF-Lac2 and PsaEF-Lac3/PsaEF-Lac4 were used for amplifying the upstream and the downstream sequences of the *psaEF* stop codon, respectively. Complementary region in primers PsaEF-Lac2 and PsaEF-Lac3 contained *Pst*I and *Sac*I restriction sites. The flanking region of the *psaEF* stop codon were fused by overlapping PCR using primers PsaEF-Lac1and PsaEF-Lac4. The resulting PCR fragments were constructed into *Eco*RI and *Hind*III sites of pUC18 to generate the plasmid pSMV35. Then, the FRT-Cm-FRT DNA fragments amplified from pKD3 was cloned into *Pst*I and *Sac*I sites of pSMV35 to generate pSMV36. pSMV37 for in-frame deletion of *psaE* was constructed by inserting the overlapping PCR fragments of the *psaE* the flanking regions into *Kpn*I and *Xma*I sites of pRE112. pSMV38 for in-frame deletion of *psaF* was constructed by inserting the overlapping PCR fragments of the *psaF* the flanking regions into *Kpn*I and *Xma*I sites of pRE112. The *psaE*-6x*his* fragment and its downstream DNA fragment were amplified by primer sets of PsaE-his1/PsaE-his2 and PsaE-his3/PsaE-his4, respectively. The overlapping fragments amplified by PsaE-his1/PsaE-his4 were cloned into *Kpn*I and *Xma*I sites of pRE112 to generate pSMV39. The *psaF-HA* fragment and its downstream DNA fragment were amplified by primer sets of PsaF-ha1/PsaF-ha2 and PsaF-ha3/PsaF-ha4, respectively. The overlapping fragments amplified by PsaF-ha1/ PsaF-ha4 were cloned into *Kpn*I and *Xma*I sites of pRE112 to generate pSMV40. The PCR fragment of *psaE*-His gene was cloned into *Bam*HI and *Hind*III sites of pUHE21–2*lacI*$^q$ to construct pSMV41. The PCR fragment of *psaF*-His gene was cloned into *Bam*HI and *Hind*III sites of pUHE21–2*lacI*$^q$ to construct pSMV42. The full promoter of *psaA* (544 base pairs in front of *psaA* start codon, $P_{psaA}1$) cloned into *Asc*I and *Kpn*I sites of pEU730, a *lacZ* promoterless plasmid, to generate pSMV43 (referred to P1). The 515 base pairs in front of *psaA* start codon was cloned into *Asc*I and *Kpn*I sites of pEU730 to generate pSMV44 (referred to P2). The 475 base pairs in front of *psaA* start codon was cloned into *Asc*I and *Kpn*I sites of pEU730 to generate pSMV45 (referred to P3). The 444 base pairs in front of *psaA* start codon was cloned into *Asc*I and *Kpn*I sites of pEU730 to generate pSMV46 (referred to P4). The 377 base pairs in front of *psaA* start codon was cloned into *Asc*I and *Kpn*I sites of pEU730 to generate pSMV47 (referred to P5). The 333 base pairs in front of *psaA* start codon was cloned into *Asc*I and *Kpn*I sites of pEU730 to generate pSMV48 (referred to P6). The 291 base pairs in front of *psaA* start codon was cloned into *Asc*I and *Kpn*I sites of pEU730 to generate pSMV49 (referred to P7). The 240 base pairs in front of *psaA* start codon was cloned into *Asc*I and *Kpn*I sites of pEU730 to generate pSMV50 (referred to P8). The 377 base pairs in front of *psaA* start codon with four-point mutations was cloned into *Pme*I and *Kpn*I sites of pEU730 to generate pSMV51 (referred to P5m). The gene fragment of *Y. pestis rovA* with its native promoter was cloned into pYA4454 to generate pSMV52.

In order to purify PsaE protein for DNase I Protection Assay, the *psaE* fused to a C-terminal 6×His was amplified from gDNA of *Y. pestis* using PsaE-His-F and PsaE-His-R and cloned into the *Nco*I and *Hin*dIII sites of plasmid pBAD-HisB to form pSMV53.

**Protein purification**

*E. coli* TOP10 carrying pSMV53 (*pasE-6xhis*) was grown overnight at 37°C in LB broth supplemented with 100 μg/ml ampicillin. The procedures for protein expression and purification were described in our previous study (Sun and Curtiss, 2012). Briefly, bacteria were grown at 37°C in a 2 L flask (agitation at 200 rpm) to an OD600 of 0.9 and then induced to production of the PsaE-6xHis protein for 3 hours through adding 0.1% arabinose. Bacteria were harvested by centrifugation at 6,000 × *g* for 10 min, resuspended in 50 ml of 50 mM sodium phosphate buffer (pH 8.0) containing 300 mM NaCl, and broken using ultra-sonication on ice. The bacterial lysate was centrifuged at 12,000 × *g* for 20 min, and the soluble fraction was applied to large SDS-PAGE gels to separate proteins. The gels containing PsaE-6xHis protein was cut and enriched via Whatman® Elutrap electroelution system (Sigma).
