## Supplementary material for "Dissecting *psa* locus regulation in *Yersinia pestis*": Table S3

**Table S3. Primers used in this work**

| Name | Sequence |
|---|---|
| psaA-lacZY1 | Cgggaattccacactaatgatcgctgcttgtggta (*Eco*RI) |
| psaA-lacZY2 | tacgagctcggcagcctgcagtcgttatttaaaatacatactcttcaac |
| psaA-lacZY3 | cgactgcaggctgccgagctcgtaatcatagaacatgatggcttgctgt |
| psaA-lacZY4 | cggaagcttccttgtaacccagcatcaagcctgaata (*Hin*dIII) |
| psaE-1 | tctgaattccagatcaggggaggtgag (*Eco*RI) |
| psaE-2 | gagagctcggcagcctgcagtctaacagcagcacctcattcttgga |
| psaE-3 | actgcaggctgccgagctctcattatctgtgatcaatcggaatg |
| psaE-4 | cggaagcttgaataggtaatacttccccgttggca (*Hin*dIII) |
| PsaE-C1 | cggggatcccagatcaggggaggtgagggcaaatg (*Bam*HI) |
| PsaE-C2 | cggaagcttctgttatagatatgagagtaagtga (*Hin*dIII) |
| psaF-1 | cggcccggggtgaggtgataaagacatataagaatg (*Xma*I) |
| psaF-2 | tgagctcggcagcctgcagcattactgttatagatatgagagtaag |
| psaF-3 | gctgcaggctgccgagctcacgtcctatcatgacgttagttaatca |
| psaF-4 | gttccggtaccggagttaatgacagtagaagcgtttgcca (*Kpn*I) |
| psaF-C1 | cggggatccgatcaatcggaatgcaaacagcaatga (*Bam*HI) |
| psaF-C2 | cggaagcttttaactaacgtcatgataggacgtatg (*Hin*dIII) |

| | |
|---|---|
| PsaEF-lacZY1 | cggggtacccagtaatgtttttcttatttttaatcta (*Kpn*I) |
| PsaEF-lacZY2 | tagagctcggcagcctgcaggataaaatgattaactaacgtcatgatag |
| PsaEF-lacZY3 | tcctgcaggctgccgagctctattgattttctattttagatgacattttta |
| PsaEF-lacZY4 | cggcccgggataactcagtcgcagacctatagatag (*Xma*I) |
| psaA P-F1 | cggggcgcgcctcattttatctattgattttctatt (*Asc*I) |
| psaA P-F2 | cggggcgcgccatgacatttttaaataataaatatggcg (*Asc*I) |
| psaA P-F3 | cggggcgcgccggtatctttagaaccgtttttactcctta (*Asc*I) |
| psaA P-F4 | cggggcgcgccgtactatgctcatcattattaatgctctttca (*Asc*I) |
| psaA P-F5 | cggggcgcgccgtcatgatagataatgaaataaaag (*Asc*I) |
| psaA P-F6 | cggggcgcgccgttttccaagattaatcttaaca (*Asc*I) |
| psaA P-F7 | cggggcgcgcccattaataattttaaacatccagaag (*Asc*I) |
| psaA P-R | cggggtaccgagaacagtctccattaaatg (*Kpn*I) |
| PpsaA-PM | cgggtttaaacgtcatgatagataatgaaataaaag |
| PsaE-His-F | cggccatgggcatgagtcactgtgttgttttaaataaattag (*Nco*I) |
| PsaE-His-R | gccaagctttcagtgatgatgatgatgatgttgctgtttgcattccgattgatcaca (*Hin*dIII) |
| psaE-6xHis-1 | cggggtaccgaatgagggctatagctatcaaaagga (*Kpn*I) |
| psaE-6xHis-2 | cagatatcagtgatgatgatgatgatgttgctgtttgcattccgattgatcacaga |
| psaE-6xHis-3 | catcatcactgatatctgtgatcaatcggaatgcaaacagcaatgaaagcaaaatcact |

| | |
|---|---|
| psaE-6xHis-4 | ccaatcccgggagcgatcctctggcca (*Xma*I) |
| psaF-HA-V1 | tgacaacctgttggcctgatcgagtcg |
| psaF-HA-V2 | tacccatacgatgttccagattacgct |
| rovA-m1 | cgggaattcattatctgcatgaatatattatcta |
| rovA-m2 | attactgcaggctgccgagctcttgctcctccttttaattagcgtg |
| rovA-m3 | caagagctcggcagcctgcagtaatttaaagttcaaagactttattca |
| rova-m4 | cggaagcttcccaggctaagcaagtacagacggtg |
| rovA-6xHis-1 | cggggtaccttggaatcgacattaggatctga |
| rovA-6xHis-2 | ttaatgatgatgatgatgatgcttagtttgtaattgaataatatttttc |
| rovA-6xHis-3 | catcatcatcatcatcattaataatttaaagttcaaagactttattcaca |
| rovA-6xHis-4 | cggcccggggatgataacaacgtagtcggtgcca |
| rovA-C1 | cggggatccagcacgctaattaaaaggaggagcaattg |
| rovA-C2 | cggggatccgaactttaaattattacttagtttgta |
